# FrustraPocket: A protein–ligand binding site predictor using energetic local frustration

**DOI:** 10.1101/2022.12.11.519349

**Authors:** Maria I. Freiberger, Camila M. Clemente, Eneko Valero, Jorge G. Pombo, Cesar O. Leonetti, Soledad Ravetti, R. Gonzalo Parra, Diego U. Ferreiro

## Abstract

Proteins are evolved polymers that minimize their free energy upon folding to their native states. Still, many folded proteins display energetic conflict between residues in various regions that can be identified as highly frustrated, and these have been shown to be related to several physiological functions. Here we show that small-ligand binding sites are typically enriched in locally frustrated interactions in the unbound state. We built a tool using a simple machine learning algorithm named FrustraPocket that combines the notion of small-molecule binding pockets and the localization of clusters of highly frustrated interactions to identify potential protein-ligand binding sites solely from the unbound forms.

**Availability and implementation (github):** https://github.com/CamilaClemente/FrustraPocket/

**Docker container:** https://hub.docker.com/r/proteinphysiologylab/frustrapocket

## Background

The Energy Landscape Theory of protein folding (1) states that natural occurring proteins are evolved systems that can robustly fold, in biological compatible times, due to the existence of a strong energetic bias towards their native states. The shape of the energy landscape of most globular proteins resembles a rough funnel where the free energy rapidly drops when interactions that are present in the native state are formed, according to the “Principle of Minimum Frustration”. However, upon folding, natural proteins may not be able to completely solve all energetic conflicts among their residues and some conflicting interactions may remain in their native states (2).

Along evolutionary times, proteins have not been optimised to fold but to function and often protein stability and function are in conflict with each other (3). In the last years, it has been shown that unresolved energetic conflicts in the native state of proteins, a.k.a highly frustrated interactions, are of great relevance to many functional aspects. Protein-protein interactions (2), allosteric sites (4), catalytic sites and co-factors binding (5), disease associated mutations (6), protein dynamics (7), disorder-to-order transitions in protein complexes (8) or evolutionary patterns in protein families (9) are some of the many functional aspects that have been related to the concept of energetic local frustration. Basically, if a protein will bind and recognise a defined substrate, we expect that there will be a set of residues near the binding site that may be in conflict with their local environment, that may become stabilised once the recognition takes place.

Identification of protein ligand binding sites (LBSs) constitutes an important step towards the elucidation of protein functions, understanding of how pathogens interact with their hosts or to grasp insights for the rational design of drugs targeting specific proteins.

In recent years, several approaches to predict LBSs have been developed. Most are based on a geometric definition of the protein’s pockets to which small-ligands bind. However, the amount of pockets that most of the tools predict is too large and their identification, based on geometric means, does not provide an intuitive way to distinguish the best candidate protein-ligand pocket.

We have developed FrustraPocket, a machine learning based algorithm that combines the notion of protein pockets and the identification of highly frustrated patches and machine learning algorithm to predict protein-ligand binding sites.

## Methods

### Dataset, local frustration and local density

To characterize the LBSs, we selected from BioLiP database (10) all entries that had their EC Number annotated in order to select only enzymatic proteins. The enzymes were classified according to their oligomeric status and a non-redundant dataset of 1007 monomeric enzyme proteins was finally selected (the PDBids used are available in GitHub CamilaClemente/FrustraPocket/).

The protein structures were downloaded from the PDB (https://www.rcsb.org/), the frustration patterns and local density (LD) were calculated using the protein FrustratometeR (11, 12) (www.frustratometer.tk).

### Pair distribution function to quantify local frustration patterns

The Mutational Frustration Index (MFI) is a measure that is assigned to the interaction between two residues (2). Therefore, to quantify the density of contacts of each frustration type (i.e highly frustrated, neutral or minimally frustrated) around a binding residue, or any residue in general, we create virtual particles (VPs) to represent the frustration assigned to the interaction between every pair of residues in contact. VPs coordinates correspond to the middle point along the euclidean distance between the interacting residues C*α* atoms. For each protein structure we obtained the list of contacts in each frustration type and calculated their VPs coordinates. Subsequently, distances from the C*α* from the selected residues, binding residues or control, or co-factor molecules were calculated with respect to the VPs coordinates. Pair distribution functions G(r) in all cases g(r) values were normalized such that g(20) = 1.

#### Feature extraction methods

##### Dataset

In order to build the dataset for the training and testing for our machine learning algorithm, we used the frustration calculation for all residues in the dataset. Furthermore, residues were classified as those that are annotated to form protein-ligand interactions (class 1) and those that do not form protein-ligand interactions (class 0). Because the amount of residues that are of protein-ligand interaction (class 1) is a very small percentage with respect to the total length of the protein, to avoid an unbalanced dataset, we used an undersampling technique. For this experiment, we have used NearMiss-1, this version selects the examples from the majority class with the smallest distance to the three closest examples from the minority class. The final dataset containing 97246 (48623 for class 1 and 48623 for class 0) amino acids and the features used are in table 1.

**Table 1.** Features used for XGBoost training and testing.

| Feature Name | Description |
| --- | --- |
| Number of contacts | Number of contacts that each residue makes |
| Local Density | Local density value of the residue |
| % of Highly Frustrated Contacts | Percentage of highly frustrated contacts that the residue makes with other residues of the residue |
| % of Highly Frustrated Contacts around of a 5 Å sphere | Percentage of highly frustrated contacts with 5 Å sphere around the $C\alpha$ |
| % of Minimally Frustrated Contacts around of a 5 Å sphere | Percentage of minimally frustrated contacts with 5 Å sphere around the $C\alpha$ |
| % of Neutral Frustrated Contacts around of a 5 Å sphere | Percentage of neutral frustrated contacts with 5 Å sphere around the $C\alpha$ |
| Class | Indicates if the amino acid is a P-L interaction residue (1) or not (0) |

##### Model construction and evaluation

Extreme Gradient Boosting (XGBoost) is a machine learning method which is widely used for data science (13). This method is a gradient boosting decision tree. The XGBoost algorithm is a Python library that implements a collection of machine learning algorithms. It provides parallel tree boosting and is one of the most important machine learning libraries for classification or regression tasks, among others. This algorithm was developed to maximize its accuracy and scalability, as well as to push the limits of computing power improving its performance and computational speed. In addition, the implementation of these models has given very good results in previous works related to this topic (14, 15) and also, because our dataset is of structured type.

The data set was randomly split into a training set (80% of the data set) and a test set (20% of the data set) using the train_test_split() function of the Sklearn library. Predictions were made on the test set. In order to check the success of the models AUC, accuracy, precision, recall, kappa and f1-score were calculated. The parameters used for XGBoost model, were,

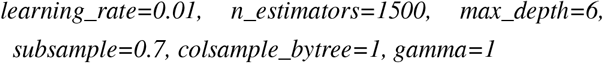

This was implemented using Python 3.6 and a Linux operating system.

## Implementation

### Algorithm workflow

FrustraPocket is available in a github repository (https://github.com/CamilaClemente/FrustraPocket/) and is coded in python3. It is also implemented in a docker container (proteinphysiologylab/frustrapocket). The workflow used to construct FrustraPocket is in Figure 1. The input of FrustraPocket should be a protein structure or a PdbID.

**Fig. 1.**
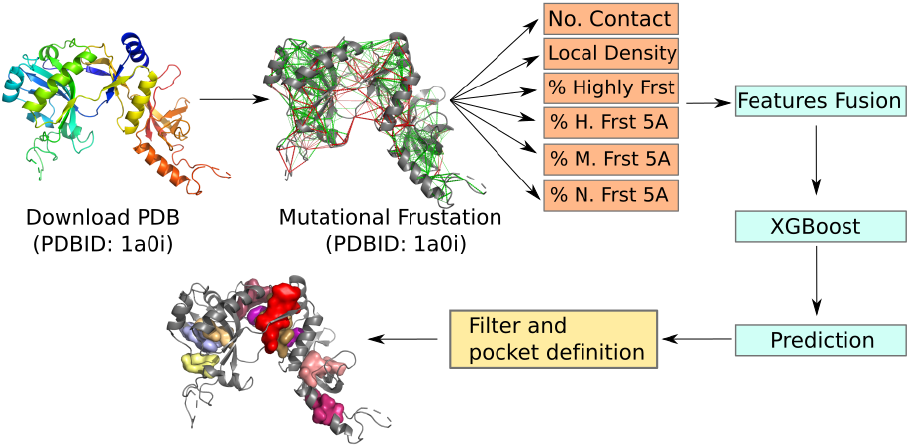
The workflow of FrustraPocket algorithm. **Step 1:** Download the PDB (only in the case that the PDB file is not provided for the user). **Step 2:** Mutational frustration calculation using FrustraR (11). **Step 3:** all the necessary features are obtained from the frustraR outputs. **Step 4:** predictions are made using the defined model. The results of the prediction are filtered and the pockets are defined.

#### Pocket predictions steps

The input file is only the PDBID or a custom PDB file generated by the user. **Step 1:** Download of the protein structure. **Step 2:** Calculation of MFI (11) and the corresponding proportion of highly frustrated interactions per residue MFI_hprop and the LD (16) of the protein. FrustratometeR calculates the percentage of the different frustration contact types (i.e highly frustrated, neutral or minimally frustrated) around a sphere of 5 Å, centered in the C*α* atom of the residue (5Adens). **Step 3:** Run the prediction using the XGBoost ML model. **Step 4:** Once the prediction is done, a pocket is defined by at least 5 close residues that are all of class 1, in case there are no class 1 residues close to each other no pocket is defined. The output files, include the frustration calculation, the pockets in PDB format, a pymol script to visualize the pockets in the protein structure and the center of mass for each pocket.

## Results

### Protein–ligand binding sites are spatially surrounded by highly frustrated interactions

To analyse the local frustration distribution in enzymatic protein–ligand binding sites, we collected all entries from the BioLiP database (10), we divided the dataset according to the oligomeric state of proteins and monomeric proteins were selected (1007 non-redundant entries). Monomeric enzymes were selected because their local frustration patterns were already analyzed (5) and we selected monomers due to their topological simplicity.Then, we calculated the local frustration patterns using the FrustratometeR package (11).

To quantify the local frustration patterns we calculated the pair distribution functions g(r) for the various classes of contacts as classified by the frustration index. The g(r) calculates the density of VPs (see methods) corresponding to the different types of contacts as a function of distance relative to the C*α* of the binding residues.

In Fig.2A we show the pair distribution function *g*(*r*) for those residues that are annotated as protein - ligand binding residues. We can see an enrichment of neutral and highly frustrated interactions, relative to the contacts topology of the protein (black lines). The distribution of interactions around the binding residues displays two characteristic peaks, one located around 1 Å, corresponding to those interactions of the binding residues themselves (first shell), and a second peak between 2 and 4 Å, which comprises interactions between residues that coordinate the binding (second shell). However, the enrichment of highly frustrated interactions in the second shell is higher than what is expected by the protein topology (black line). Note that in both first and second shells there is a depletion of minimally frustrated interactions. These results show that the specific sites for protein-ligand recognition are typically frustrated in the unbound state.

**Fig. 2.**
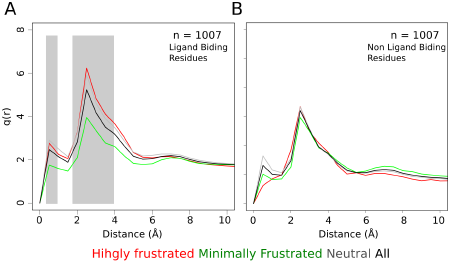
Pair distribution functions, g(r), between the C*α* of a) the annotated binding residues and the center of mass of the contacts and b) control residues. Green, minimally frustrated contacts; red, highly frustrated contacts; gray, neutral contacts; black, all contacts. g(r) plots were adjusted in their axis ranges to enhance visualizations; however, in all cases g(r) values were normalized such that g(20) = 1. **A** Residues that are annotated as protein - ligand binding residues. **B** Control residues, defined as randomly selected residues that are no annotated as protein - ligand binding residues. In gray are shown the 1st shell and the 2nd shell, respectively.

In Fig.2B, we observe the *g*(*r*) for a control set that was generated using random residues that are not involved in binding sites that shows no enrichment, in contrast to what is observed for ligand binfing residues Fig.2A. Based on this and in previous work where we observed an enrichment of highly frustrated interactions around protein-ligand interaction residues and also at catalytic sites (5) we decided to exploit this feature and use it to predict protein-ligand interaction and catalytic sites by combining information of the local frustration patterns and the local density of residues in a protein structure.

### Performance of the model

In order to the evaluate the effectiveness of the XGBoost model implemented in this work, we used the Receiver Operating Characteristic (ROC) curve (Fig. 3). The True Positive rate is defined as TP/(TP+FN) and the False Postive rate is defined as FN/(FN+TP), where TP is the True Positive and FN is the False Negative. The values for accuracy, precision, recall, kappa, f1-score are provided in Table 2. The AUC obtained by XGBoost is 0.70, indicating that our method does not select residues randomly. These values indicate that our model correctly detects approximately 70% of the ligand-protein binding residues, solely based on their local frustration patterns in the unbound states.

**Fig. 3.**
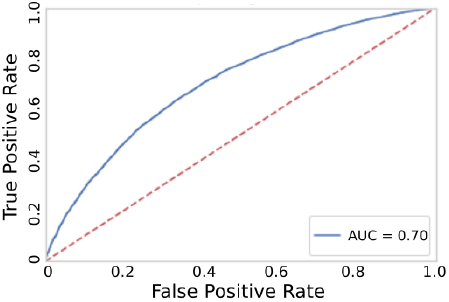
The ROC curve of XGBoost applied to the local frustration patterns of monomeric enzymes. The true negative rate is the probability that an actual positive will test positive and the false negative rate is the probability that a true positive will be missed by the test

**Table 2.** Metrics used to evaluate the performance of the model.

| Metric | Value |
| --- | --- |
| AUC | 0.70 |
| Accuracy | 0.64 |
| Precision | 0.64 |
| Recall | 0.66 |
| Kappa | 0.29 |
| f1-score | 0.65 |

#### Particular Example

As an example, we have applied FrustraPocket to the ATP-dependent DNA ligase (PdbID: 1A0I). Fig.4A-B represents the first step of the tool where Fig.4A shows the mutational frustration frustratogram and Fig.4B shows the LD of each residue of the protein. Fig.4C represents the output of the nine predicted pockets of the protein and fig.4D the center of mass for each predicted pocket.

**Fig. 4.**
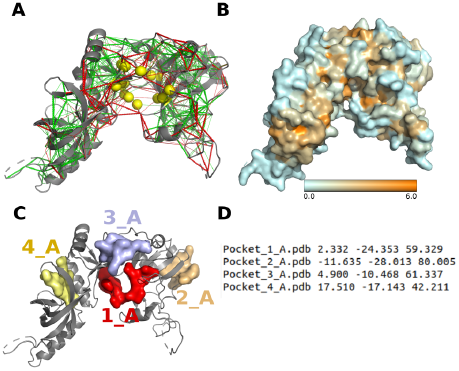
Example PdbID: 1A0I. **A.** The backbones of the proteins are shown as gray cartoons, minimally frustrated contacts are depicted with green lines, highly frustrated interactions with red lines. Neutral interactions were omitted to help interpretation. **B.** Residues with lower local density are shown in blue and residues with higher local density are shown in orange. **C.** Output of the tool, in gray the crystallised ligand, in different colors the predicted pockets. **D.** Center of mass for each pocket.

## Discussion and conclusion

The identification of small-ligand binding sites in a protein structure is a central theme to protein physiology and has been the subject of an increasing number of studies in the last decade. Currently there are many predictors, most of them based on geometry and until now there was no method based on the analysis of the protein only energetics. Energetic local frustration is a biophysics-based concept related to various functional aspects of proteins (2, 4–9), especially to the interactions between proteins or proteins and their ligands. Hence, we consider that frustration is an intuitive concept that can improve prediction of LBSs as we have shown in this article.

Here we calculated and characterize the frustration patterns for monomeric enzymes (n = 1007) that have their LBSs annotated in BioLiP database (10). We have found that the residues that are implicated in protein - ligand interactions are enriched in highly frustrated interactions.

Thus, Frustrapocket may be a valuable tool to identify unknown potential ligand-binding sites. Indeed, the fact that in most cases we identify pockets for which no ligand is known may be hinting that several cryptic binding sites are present in many proteins.

## ACKNOWLEDGEMENTS

This work was supported by the Consejo de Investigaciones Cientificas y Tecnicas (CONICET), the Universidad de Buenos Aires (UBACYT 2018-20020170100540BA, Grant PICT2016/1467 to D.U.F.). D.U.F and S.R are a CON-ICET researchers and M.I.F. and C.M.C are holders a CONICET fellowship.

